# Functional specialization of the subdomains of a bactofilin driving stalk morphogenesis in *Asticcacaulis biprosthecum*

**DOI:** 10.1101/2024.12.16.628611

**Authors:** Maxime Jacq, Paul D. Caccamo, Yves V. Brun

## Abstract

Bactofilins are a recently discovered class of cytoskeletal protein, widely implicated in subcellular organization and morphogenesis in bacteria and archaea. Several lines of evidence suggest that bactofilins polymerize into filaments using a central β-helical core domain, flanked by variable N- and C-terminal domains that may be important for scaffolding and other functions. However, a systematic exploration of the characteristics of these domains has yet to be performed. In *Asticcacaulis biprosthecum*, the bactofilin BacA serves as a topological organizer of stalk synthesis, localizing to the stalk base and coordinating the synthesis of these long, thin extensions of the cell envelope. The easily distinguishable phenotypes of wild-type *A. biprosthecum* stalks and *ΔbacA* “pseudostalks” make this an ideal system for investigating how mutations in BacA affect its functions in morphogenesis. Here, we redefine the core domain of *A. biprosthecum* BacA using various bioinformatics and biochemical approaches to precisely delimit the N- and C-terminal domains. We then show that loss of these terminal domains leads to cells with severe morphological abnormalities, typically presenting a pseudostalk phenotype. BacA mutants lacking the N- and C-terminal domains also exhibit localization defects, implying that the terminal domains of BacA may be involved in its subcellular positioning, whether through membrane interactions through the N-terminal domain or through interactions with the stalk-specific morphological regulator SpmX through the C-terminal domain. We further show that point mutations that render BacA defective for polymerization lead to stalk synthesis defects. Overall, our study suggests that BacA’s polymerization, membrane association, and interactions with other morphological factors all play a crucial role in the protein’s function as a morphogenic regulator. The specialization and modularity of the terminal domains may underlie the remarkable functional versatility of the bactofilins in different species.

**Author summary:** Bacteria exhibit a wide variety of shapes and structures, many of which are crucial for their cellular functions. Among these structures is the stalk—a thin, tubular extension of the cell envelope formed by bacteria such as *Asticcacaulis biprosthecum*. Stalk synthesis in *Asticcacaulis biprosthecum* relies on the bactofilin BacA, a self-polymerizing cytoskeletal protein, whose deletion results in the dysregulation of stalk synthesis, and the formation of short, stubby “pseudostalks”. We use this unique phenotype to characterize the subdomains of BacA, and find that BacA’s ability to coordinate stalk synthesis depends on its conserved polymerization domain as well as its flanking N- and C-terminal domains, which are essential for proper localization and interactions. Our findings highlight how bactofilins combine conserved and variable regions to generate complex structures that serve as a platform for evolving new functions.

## Introduction

All cells require spatiotemporal organization of their components and processes for proper function. While bacteria lack the typical membrane-bound organelles that allow for higher-order partitioning in eukaryotic systems, they possess cytoskeletal scaffolding proteins for spatiotemporal organization of protein complexes. Bacterial cytoskeletal scaffolds facilitate the localization and interplay of proteins and their regulatory elements at specific subcellular locations at specific times, partitioning the cytoplasm into functionally distinct domains. Given this role, bacterial cytoskeletal scaffolds serve as key regulators of many important processes in bacterial cells, including cell growth, cell division, DNA segregation, establishment of cell polarity, morphology, and magnetotaxis (1,2).

The most studied bacterial cytoskeletal scaffolds are MreB (3), FtsZ (4), and Crescentin (5), which represent large families of prokaryotic homologs of actin, tubulin, and intermediate filaments, respectively. However, bacteria also possess more diverse cytoskeletal scaffolds (6). Among these are the bactofilins, a recently discovered class of cytoskeletal proteins that are widely distributed throughout bacteria and archaea, and occasionally found in a few eukaryotes (7–11). Bactofilin proteins are characterized by a core bactofilin domain (DUF583) of ∼100 amino acids, flanked by variable N-and C-terminal regions (8–10,12–14). Like other bacterial cytoskeletal proteins, bactofilins are known to polymerize. Nonetheless, little is known about how the specific domains and polymerization properties of bactofilins affect their functions *in vivo*.

Functionally, bactofilins are often implicated in cellular morphogenesis (15–18), likely through their mediating role in the synthesis of the peptidoglycan (PG) cell wall, a bacterial exoskeleton that gives cells their shape. In *Proteus mirabilis*, deletion or truncation of the bactofilin *ccmA* results in highly curved, morphologically defective cells (16). In *Myxococcus xanthus*, the bactofilin BacM assembles into fibers throughout the cell, and deletion of *bacM* results in crooked-rod shaped cells that show increased sensitivity to antibiotics targeting the cell wall (18,19). In *Helicobacter pylori, ccmA* deletion leads to loss of the cell’s characteristic helical shape, and truncation of the N-terminal region of CcmA results in a phenotype similar to *ΔccmA* (15,20,21).

*A. H. pylori* CcmA interacts with various morphological proteins such as Csd5 and Csd7, using its bactofilin domain (15,20). Among these, Csd7 is known to be important for the stability of a PG hydrolase called Csd1, further supporting a role for bactofilins in the synthesis and remodeling of the PG cell wall (20).

Bactofilins are also involved in stalk synthesis in the Alphaproteobacteria (9,22–24). Stalks are thin, cylindrical projections of the cell envelope, whose synthesis requires a specialized form of zonal PG synthesis in a spatially constrained zone from which the stalk is extruded (25–28). In *Caulobacter crescentus,* the bactofilins BacA and BacB contribute to the elongation of a single, polar stalk, interacting with the PG synthase PbpC at the stalked pole (9,12). Although BacA and BacB are not essential for stalk synthesis in *C. crescentus*, their deletion leads to a reduction in stalk length. On the other hand, in the related stalked species *Asticcacaulis biprosthecum*, which has bilateral stalks at the midcell, the bactofilin BacA serves as an important topological organizer of stalk synthesis - it localizes at the base of each stalk, where it anchors the developmental regulator of stalk synthesis SpmX, which in turn regulates PG synthesis at the base of the stalk. In the absence of BacA, *A. biprosthecum* cells produce shorter and wider protrusions in place of the stalk. These “pseudostalks” are characterized by unregulated PG insertion, following mislocalization of SpmX throughout the pseudostalk (29). Interestingly, in certain Alphaproteobacteria, stalks are used for reproduction, budding off daughter cells from their tips (30,31). In one such budding bacterium, *Hyphomonas neptunium*, deletion of the bactofilin genes leads to multiple, irregularly shaped, misplaced stalks, which later produce amorphous bulges corresponding to unconstrained PG insertion during bud formation (23). Thus, the bactofilins play an important role in regulating zonal growth for stalk synthesis, potentially through interactions with stalk-specific developmental regulators and PG synthetic complexes.

The bactofilins’ ability to polymerize appears to be important for their function. Polymerization is thought to be mediated via hydrophobic interactions between conserved hydrophobic residues in the core bactofilin domain, without any needed co-factors (12). Point mutations in *C. crescentus bacA* have identified residues that disrupt polymerization *in vivo*, which are important for its localization to the stalked pole in cells (12). Analogous mutations in *H. pylori* CcmA also disrupt polymerization *in vitro*, result in mislocalization *in vivo,* and produce a morphology indistinguishable from the *ΔccmA* mutant (32). Taken together, these results show that bactofilin polymerization can be easily disrupted by mutating selected residues, and that polymerization is required for the proper localization and function of bactofilins *in vivo*.

Structural analyses of the bactofilin TtBac from *Thermus thermophilus* have further elucidated the molecular interactions involved in bactofilin polymerization. TtBac protofilaments polymerize through a β-stacking mechanism similar to amyloid fibers. Interestingly, the monomers in the protofilament are organized in a non-polar fashion, with alternating N-terminal to N-terminal and C-terminal to C-terminal interactions (8). The N-terminal tails are thought to mediate inter-protofilament interactions, allowing for the formation of protofilament bundles. Furthermore, a short hydrophobic motif in the N-terminal domain appears to mediate bactofilin binding to lipid membranes *in vitro* and *in situ* (8), consistent with the bactofilins’ ability to recruit proteins to specific sites in the cell envelope *in vivo*.

The domain architecture and polymerization properties of the bactofilins are thought to be highly conserved among bacteria, but it is unclear how they contribute to bactofilin function *in vivo*. Here, we utilize *A. biprosthecum* as a morphogenic system to examine this question, applying mutations that affect its bactofilin BacA’s polymerization and N- and C-terminal domain function.

*A. biprosthecum* is an ideal *in vivo* system to examine the functional effects of such mutations, as wild-type stalks and *ΔbacA* pseudostalks provide easily distinguishable phenotypes. We present data that show that polymerization, as well as the disordered, proline-rich N- and C-terminal domains of BacA, are required for its function; mutations abolishing these features lead to severe defects in stalk synthesis, ranging from fewer and shorter stalks, to full pseudostalk synthesis. Next, we show that these mutations affect the localization dynamics of BacA. These results, along with other disparate lines of evidence, suggest that bactofilins represent an evolutionary Swiss army knife, a common platform (the self-polymerizing central β-helical core domain) upon which various adaptations can be made (using the N- and C-terminal domains) to address different biological functions.

## Results

### Defining the core bactofilin domain and the flanking regions in A. biprosthecum BacA

Bactofilin proteins can self-polymerize, which is important for their function in different bacteria (10). Bactofilin polymerization is solely dependent on the conserved central bactofilin domain of the protein, whereas the flanking N- and C-terminal domains often vary between genera and species. To gain insight into the domain architecture of *A. biprosthecum* BacA (Abi_34180; hereafter referred to as BacA), we first wanted to precisely map its domains in comparison to *C. crescentus* and other related species in the family *Caulobacteraceae*.

The predicted bactofilin domain (DUF583) of BacA, as determined by bioinformatic Pfam analysis, spans residues 52 to 138 of the protein. However, this was shorter than the overall sequence conservation observed in the central domain of this protein among closely related species within the Caulobacteraceae family (Fig. 1A). Indeed, the Pfam-predicted bactofilin domain in this family excluded several highly conserved amino acids. Moreover, bactofilin domains are usually flanked by proline rich N- and C-terminal domains (9), but none of the conserved residues adjacent to the Pfam-predicted bactofilin domains were prolines (Fig. S1A). An examination of BacA with disorder prediction software revealed that residues 39 to 145 were likely structured, which includes several conserved residues excluded from the Pfam-predicted bactofilin domain (Fig. 1B). Upon conducting an AlphaFold structure prediction of BacA and juxtaposing it with the experimentally determined protein structure of *C. crescentus* BacA (_Cc_BacA), it became evident that the current annotation of the bactofilin domain (DUF583) by Pfam falls short of capturing the complete structured bactofilin domain of BacA (Fig. 1C, S1B). Notably, the Pfam-predicted bactofilin domain lacks an entire helix at the N-terminal end and the last few amino acids at the C-terminal end of the bactofilin domain (Fig. 1C, left). Based on these observations, we predicted that the bactofilin domain of BacA is composed of amino acid (aa) residues 44 to 140, which match the ordered β-helical core of _Cc_BacA. However, several residues following 140Q exhibit notable conservation among the *Caulobacteraceae*. Consequently, we opted to assess if a bactofilin domain missing these residues would still be able to polymerize properly. To test this, we overexpressed and purified the bactofilin domain encompassing residues 44-140 (hereafter referred to as BacA_ΔNΔC_) in *E. coli* and found that it demonstrated successful filament formation *in vitro* (Fig. 1D and S1C). This demonstrates that the conserved residues beyond 140Q are unnecessary for polymerization and can thus be excluded from our updated definition of the BacA bactofilin domain. Notably, a shorter stretch of the BacA bactofilin domain (aa 44-138) failed to form filaments and precipitated in solution without any filamentation observed (S1D).

**Figure 1.**
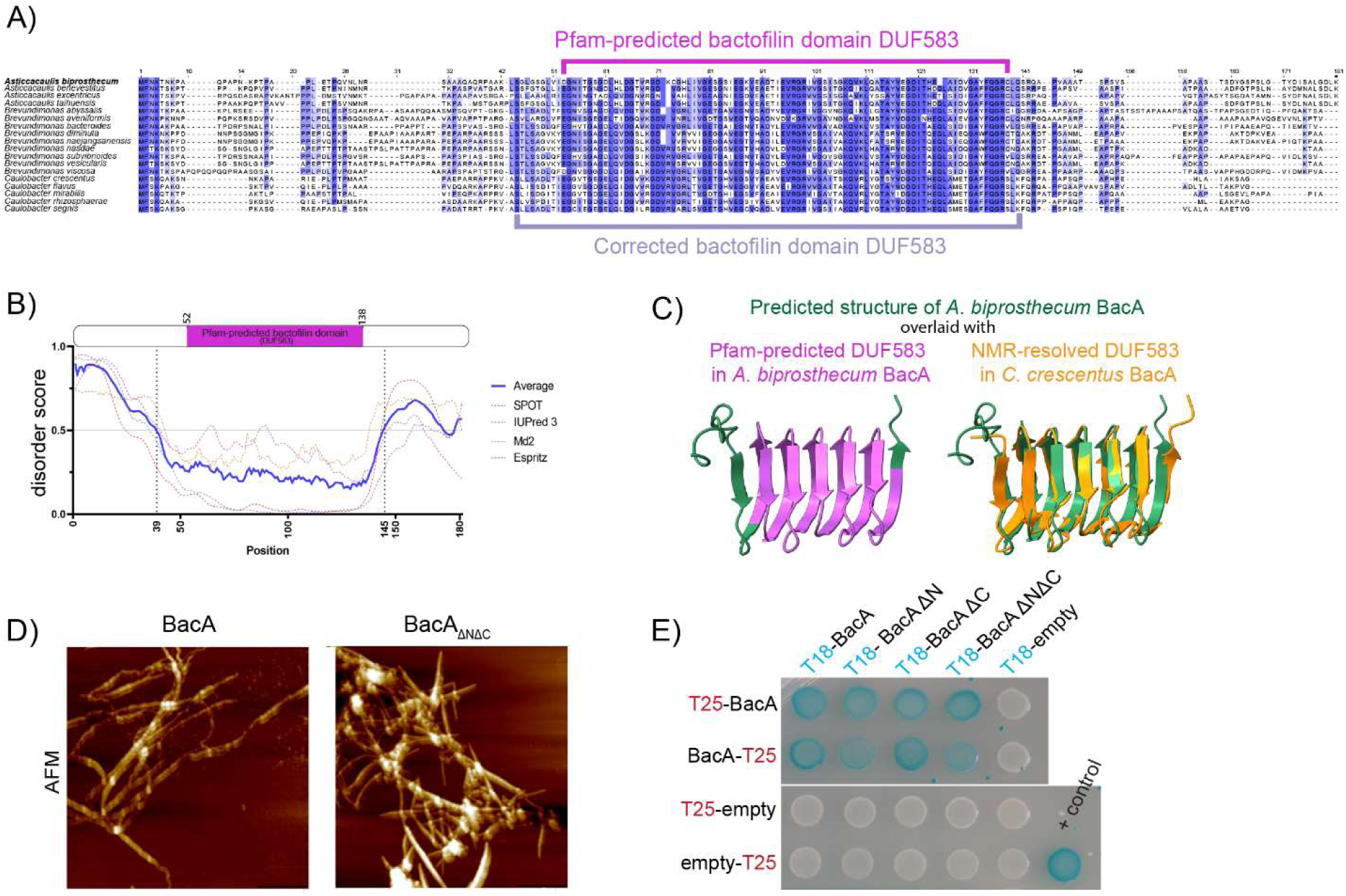
The bactofilin domain of *A. biprosthecum* BacA extends beyond the Pfam-predicted domain sequence, and forms filaments *in vitro* even in the absence of the BacA N- and C-terminal domains. **A)** Multiple sequence alignment of bactofilin sequences from various species from the *Caulobacteraceae* family. The sequences are colored using a percentage-identity color scheme, which color-codes based on conservation (dark blue: above 80%; blue: above 60%; light blue: above 40%). The Pfam-predicted bactofilin domain (DUF583) is indicated with a magenta bracket, and the corrected bactofilin domain with a mauve bracket. **B)** Disorder plot of the *A. biprosthecum* BacA sequence, generated using SPOT-Disorder2, IUPred 3, Md2, and Espritz. Disorder scores were normalized from 0 to 1. Amino acid positions with a disorder score above 0.5 are considered disordered regions of the protein. A schematic of the BacA protein indicating its Pfam-predicted bactofilin domain (DUF583), is aligned with the disorder plot as reference. **C)** Left: Structural superposition of the structure of *A. biprosthecum* BacA predicted using AlphaFold (green), with its Pfam-delimited bactofilin domain (DUF583) in magenta. Right: Structural superposition of the predicted structure of *A. biprosthecum* BacA (green) with the NMR-resolved structure of *C. crescentus* BacA (PDB-ID: 2N3D, orange). **D)** High-resolution AFM images of purified BacA filaments and BacA_ΔNΔC_. Scale bar = 1 µm. **E)** Bacterial two-hybrid (BACTH) assay in which pairs of proteins are fused to T18 and T25 fragments of adenylate cyclase and co-expressed in *E. coli*. Protein pairs that interact *in situ* enable the T18 and T25 fragments to carry out adenylate cyclase activity, which is reported as blue patches on beta-galactosidase culture plates. Here, T18-BacA constructs, with or without the BacA N- and C-terminal domains, are probed against full-length BacA fused to the T25 probe either on its N- or C-terminal. A strain co-expressing T18 and T25 fragments fused to interacting segments of a Leucine Zipper motif was used as a positive control (blue patch, bottom right). Negative controls (white patches) were performed against unfused T25 fragments.

In contrast to the structured bactofilin domain which drives polymerization, the N- and C-terminal domains of bactofilins are predicted to be highly disordered, and their functions are less understood. To investigate their importance, we designed truncation mutations of BacA, annotated as BacA_ΔN_ (Δ2-43), BacA_ΔC_ (Δ141-181), and BacA_ΔNΔC_ (Δ2-43 and Δ141-181), removing either the N- or C-terminal domains or both (Fig. S1E). In all cases, the truncated BacA proteins were able to interact with wild-type (WT) BacA in a bacterial adenylate cyclase two-hybrid (BACTH) assay in *E. coli* (Fig 1E).

### Role of the N- and C-terminal domains of BacA in stalk synthesis

To examine if BacA’s non-polymerizing regions are important for stalk synthesis in *A. biprosthecum*, we created BacA mutants lacking the N- and C-terminal domains, expressed at the native *bacA* locus in *A. biprosthecum*, with an mVenus tag to observe their localization and assess their expression (Fig. S2A). Full length BacA-mVenus expressed from the native *bacA* locus was used for comparison, to determine the function of these truncated mutants. Deletion of the entire *bacA* gene or BacA’s N- or C-terminal domains resulted in a pseudostalk phenotype, with most stalked cells producing misshaped protrusions at the site of stalk synthesis (Fig 2A-B). The *bacAΔN-mVenus* strain demonstrated a phenotype nearly as severe as *ΔbacA,* with only 5 ± 1% of these cells forming WT-like stalks, compared to 46 ± 3% in the *bacA-mVenus* strain (Fig. 2C-D). Instead, most stalked cells of the *bacAΔN-mVenus* strain formed enlarged, short pseudostalks similar in size (1.5 ± 0.8 μm long) and diameter (267 ±92 nm wide) to *ΔbacA* pseudostalks (Fig. 2E-F). Thus, loss of the N-terminal domain of BacA nearly completely abolishes BacA function *in vivo* in terms of regulating stalk synthesis, despite its ability to interact with WT BacA in BACTH experiments, and the retained ability of BacA_ΔNΔC_ to self-interact *in vitro* (Fig. 1D-E). In contrast, absence of the C-terminal domain of BacA led to a less severe phenotype. A greater proportion of *bacAΔC-mVenus* mutants had WT-like stalks (19 ± 4%), while other stalked cells produced pseudostalks with phenotypes intermediate to WT and Δ*bacA* (Fig 2A-F). The vast majority of WT-like stalks in BacA_ΔC_-mVenus cells emerged from a pseudostalk base, from which extruded a prolonged, thin stalk (Fig. 2A-B and S2B). This suggests that loss of the C-terminal domain of BacA is detrimental to stalk formation, but nonetheless allows partial functionality of the BacA_ΔC_ protein.

**Figure 2.**
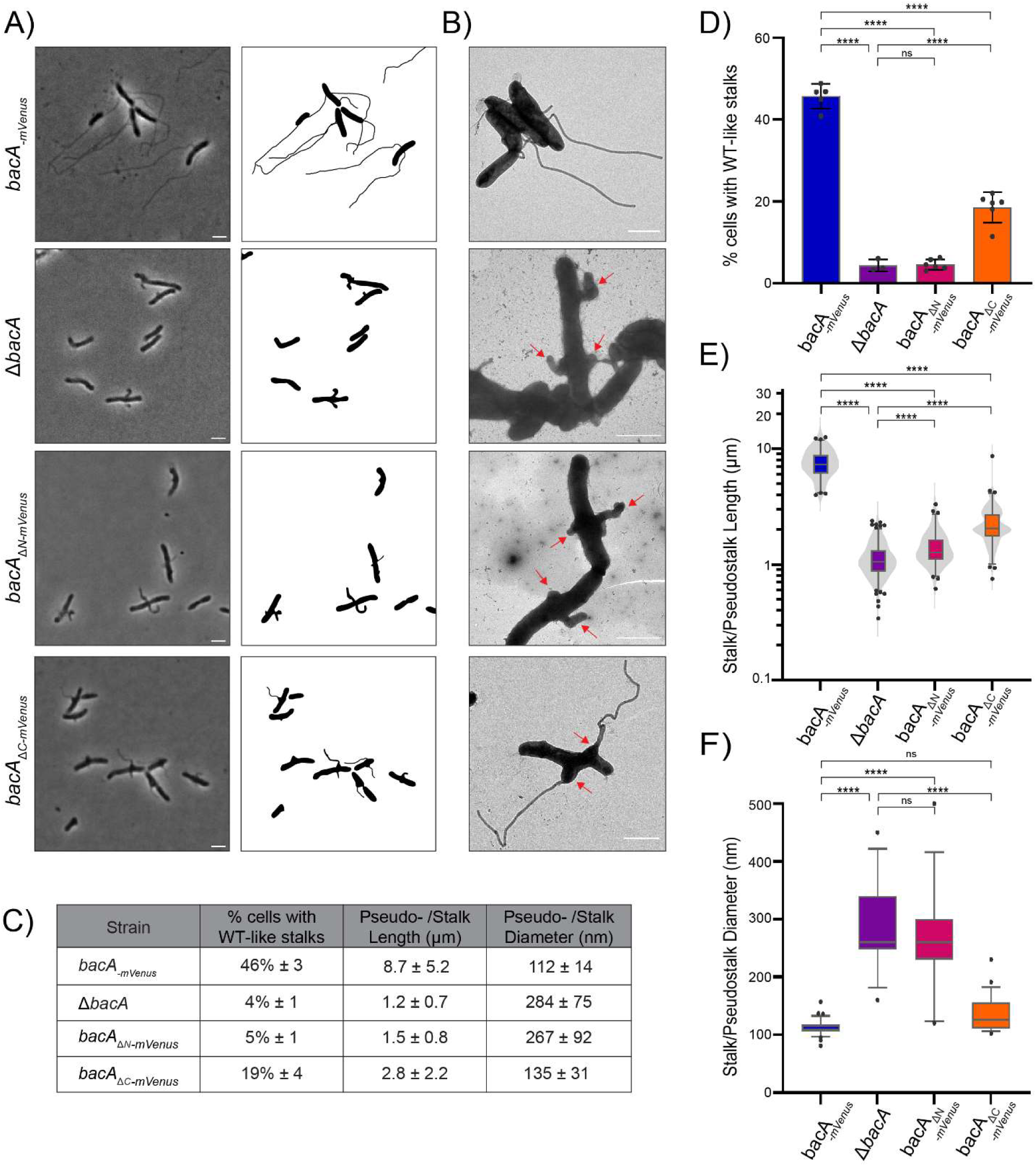
The disordered N- and C-terminal domains of BacA are required for normal morphogenesis of the stalk. **A)** Phase contrast images (left) and corresponding cell silhouettes (right) of *A. biprosthecum bacA-mVenus*, Δ*bacA*, *bacA*_Δ_*N-mVenus* and *bacA*_Δ_*C-mVenus,* to analyze stalk morphology. Cells were grown in phosphate-limited (HIGG) medium (see Methods). Scale bars = 2 μm. **B)** Transmission electron microscopy of the same strains as in Panel A. Pseudostalks are indicated with red arrows. Scale bars = 2 μm. **C)** Summary statistics for data presented in Figures 2D-F. Data shown as mean ± SD. **D)** Percentage of cells with WT-like stalks. Based on phase images, cells were scored as having a WT-like stalk (i.e. a thin extension from the cell body). Cells exhibiting thick and aberrant pseudostalks were excluded (*bacA-mVenus* n=3966; Δ*bacA* n=2749; *bacA*_Δ_*N-mVenus* n=3034; *bacA*_Δ_*C-mVenus* n=3879). Data are presented as the mean percentage of cells with WT-like stalks. Error bars represent SD. (****p ≤ 0.001, ns = not significant p > 0.05; t-test) **E)** Distribution of stalk/pseudostalk lengths in the different strains (*bacA-mVenus* n=575; Δ*bacA* n=776; *bacA*_Δ_*N-mVenus* n=448; *bacA*_Δ_*C-mVenus* n=576). Data are represented as box and whisker plots and violin plots. (****p ≤ 0.001; t-test) **F)** Distribution of stalk/pseudostalk base diameter in the different strains based on atomic force microscopy images (*bacA-mVenus* n=32; Δ*bacA* n=13; *bacA*_Δ_*N-mVenus* n=15; *bacA*_Δ_*C-mVenus* n=23; see Fig. S2B for representative images). Data are represented as box and whisker plots plots. (****p ≤ 0.001, ns = not significant p > 0.05; t-test)

Next, we examined the localization of the tagged BacA mutant constructs. Whereas WT BacA-mVenus localized as two discrete, bilateral foci at the midcell, consistent with the sites of stalk synthesis, BacA_ΔN_-mVenus no longer showed bilateral foci (Fig. 3A). Instead, BacA_ΔN_-mVenus foci were weaker and could be observed throughout the cell except at the poles, sometimes falling near the site of pseudostalk synthesis. BacA_ΔC_-mVenus showed a more intermediate phenotype, localizing laterally at the membrane, but forming elongated foci that may correspond to BacA_ΔC_-mVenus filaments (red arrowheads, Fig. 3A), perhaps from an accumulation of the protein due to its failure to consistently localize to the stalk synthesis site. Nevertheless, whenever a WT-like stalk or an intermediate stalk phenotype was observed in this strain, BacA_ΔC_-mVenus foci were always present at or near the base of these stalks.

**Figure 3.**
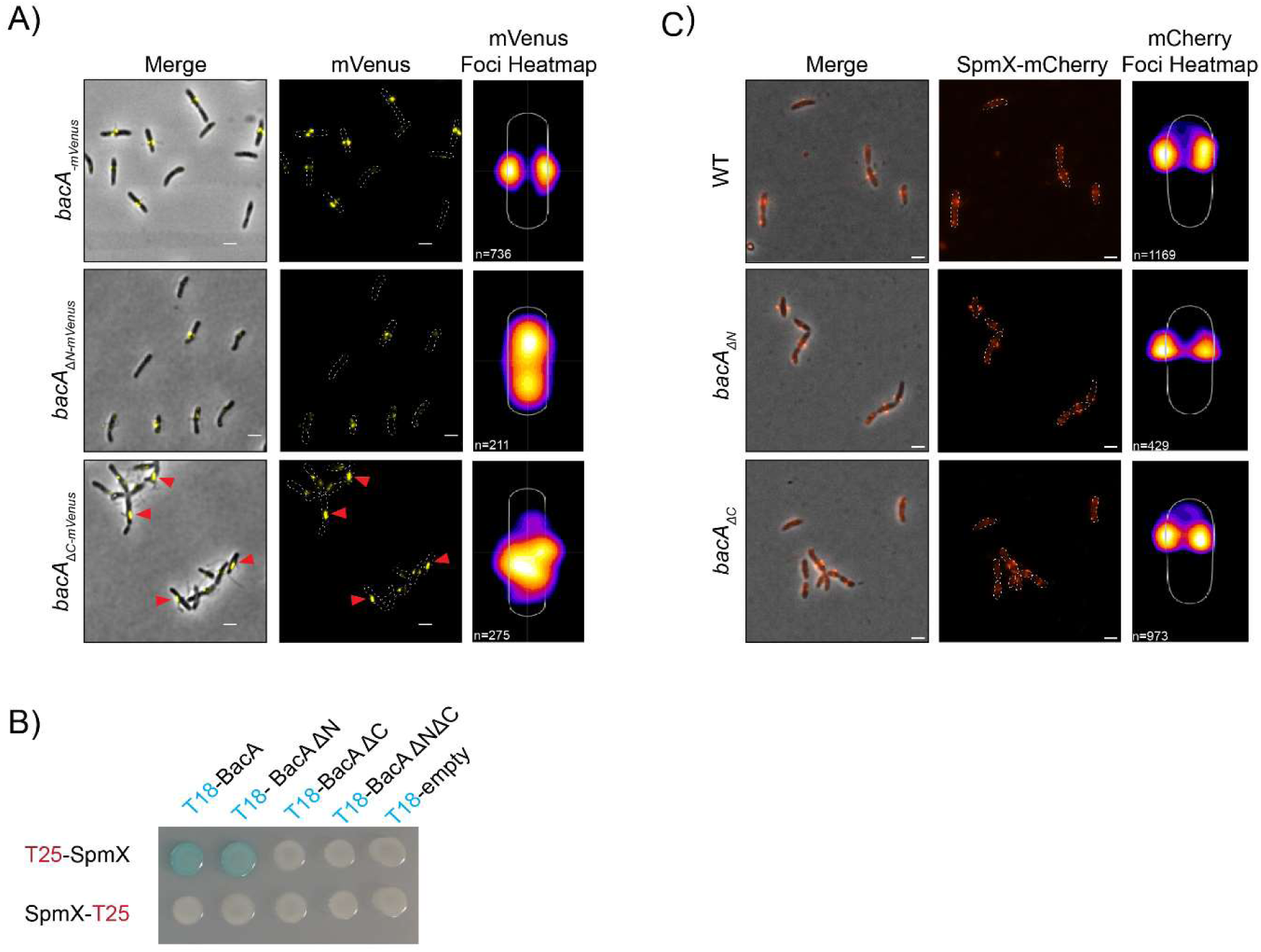
The N- and C-terminal domains of BacA are necessary to localize BacA at the site of stalk synthesis but are not necessary for SpmX localization despite impaired interaction. **A)** Merged phase contrast/fluorescence (left) and fluorescence (middle) microscopy images with localization heatmaps (right) of *A. biprosthecum* WT BacA-mVenus, BacA_ΔN_-mVenus, and BacA_ΔC_-mVenus. Red arrowheads indicate elongated foci of BacA_ΔC_-mVenus. The number of cells analyzed is shown on the bottom left of each heatmap. Scale bars = 2 μm. **B)** Bacterial two-hybrid (BACTH) assays as presented in Figure 1E, with T18-BacA constructs with or without the BacA N- and C-terminal domains, probed against SpmX fused to the T25 probe either on its N- or C-terminal. **C)** Merged phase contrast/fluorescence (left) and fluorescence (middle) microscopy images with localization heatmaps (right) of SpmX-mCherry in *A. biprosthecum* WT, *bacAΔN*, and *bacAΔC* strains. The number of cells analyzed in each case is shown on the bottom left of each heatmap. Scale bars = 2 μm

In *A. biprosthecum,* the developmental regulator of stalk synthesis, SpmX, is required for localization of BacA to the base of the stalk. Conversely, BacA functions to stabilize SpmX at the stalk base, and the two proteins interact directly (29). To test whether the N- or C-terminal domains of BacA are involved in interactions with SpmX, we conducted a BACTH assay in *E. coli* and found that loss of the C-terminal domain of BacA led to a loss of SpmX-BacA interactions, while deletion of the N-terminal domain did not (Fig. 3B). We also tested whether SpmX localization was impacted in untagged *bacA* mutant strains, by expressing SpmX-mCherry in these backgrounds. We found that SpmX-mCherry localization was not impaired in *bacAΔN* or *bacAΔC* (Fig. 3C), in both of which it localized at the stalk base as in WT cells, despite a small increase in cytoplasmic fluorescence in the *bacAΔC* mutant.

### Role of bactofilin polymerization in stalk synthesis

To assess the impact of bactofilin polymerization on stalk formation, we aimed to identify key residues at both the N-to-N and C-to-C polymerization interfaces of the core domain. We referred to previous studies elucidating the effects of point mutations on bactofilin polymerization in other species (8,12,33), and derived insights from sequence conservation among BacA homologs as well as from structural predictions of the *A. biprosthecum* bactofilin domain (Fig. 4A-B). In other species, the most impactful substitution disrupting higher-order polymerization was in the C-to-C interface of the bactofilin domain: F130A in *C. crescentus*, L110S in *H. pylori*, and F105R in *T. thermophilus* (8,12). All of these residues align with *A. biprosthecum* F134, which we substituted with an arginine for testing (Fig. 4A-B). When we overexpressed and purified the BacA_F134R_ mutant protein in *E. coli,* we were unable to detect filament formation or higher-order structures via electron microscopy. Size-exclusion chromatography data showed that purified BacA_F134R_ eluted in a single peak at around 14 mL, corresponding to a globular protein of 45 kDa, which corresponds to the size of a BacA dimer (blue line, Fig. 4C [i]). Since bactofilin filaments are known to polymerize in a non-polar manner (i.e. alternating N-to-N and C-to-C interactions), our data suggest that the BacA_F134R_ mutation may disrupt C-to-C interactions, preventing multimerization, while preserving N-to-N interactions, allowing dimerization.

**Figure 4.**
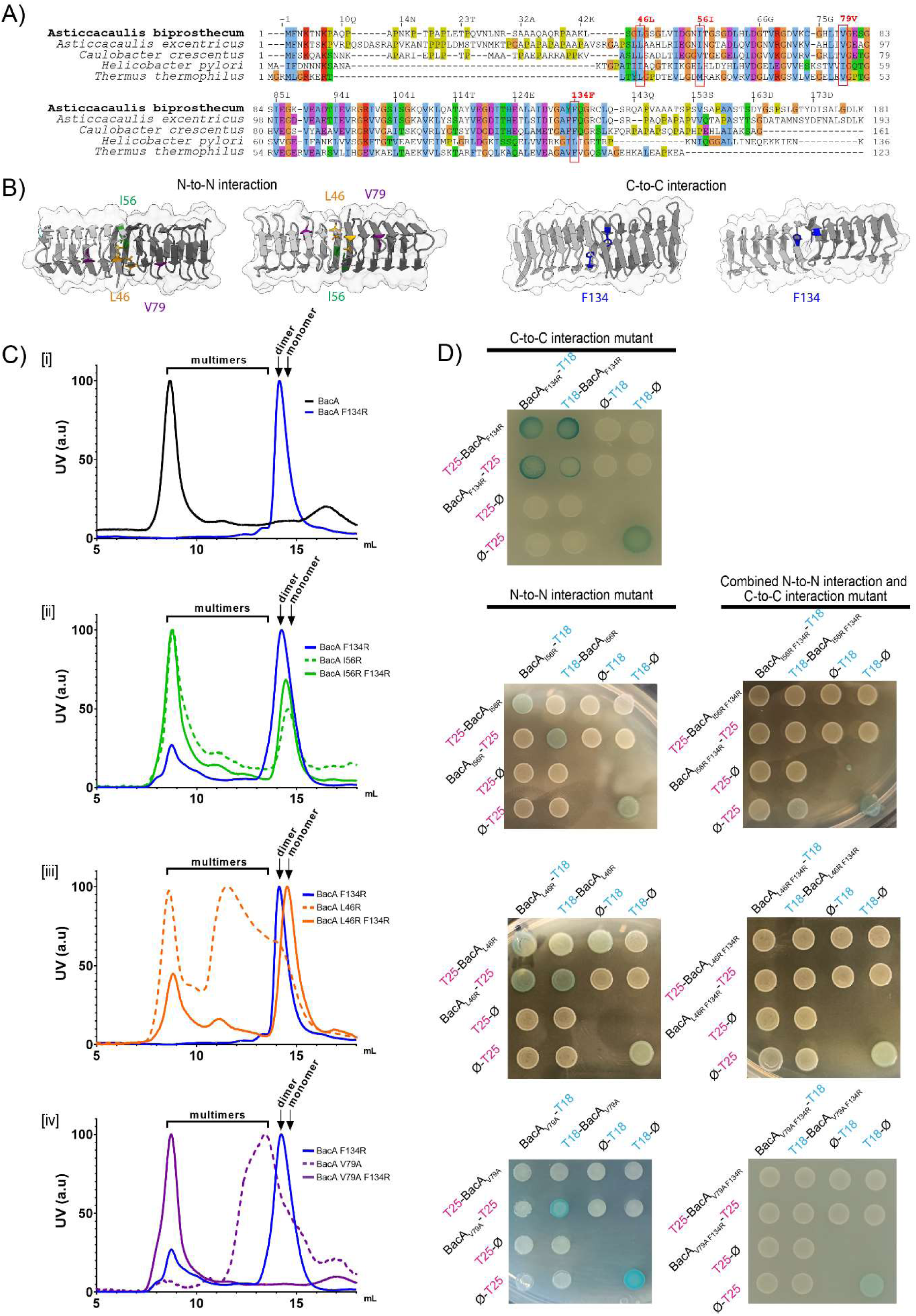
Various amino acid residues in the N-to-N and C-to-C interfaces are implicated in BacA polymerization. **A)** Multiple sequence alignment of bactofilin sequences from *A. biprosthecum*, *A. excentricus*, *C. crescentus, H. pylori*, and *T. thermophilus*. The alignment is colored using the Clustal X color scheme, which color codes amino acid residues with shared properties. The red boxes indicate conserved amino acids that have been shown to be important for bactofilin polymerization in other species, selected for substitution in *A. biprosthecum* in this study. **B)** Structural representation of the BacA dimer derived from the BacA tetramer structure predicted using AlphaFold, showing either the N-to-N interface (left) or the C-to-C interface (right). Mutated residues are highlighted. **C)** Analytical size exclusion chromatography of purified BacA carrying various mutations. Elution profiles are color coded as follows - Black: WT BacA; Blue: BacA_F134R_; Green: BacA_I56R_; Orange: BacA_L46R_; Purple: BacA_V79R._ Single N-to-N interface mutants are displayed with dashed lines while double F134R and N-to-N interface mutants of BacA are displayed as solid lines. Subpanels [i] to [iv] highlight different constructs: [i] F134R; [ii] I56R, [iii] L46R, [iv] V79A. Curves are normalized such that the maximum intensity peak is 100. The expected elution ranges of monomers, dimers and multimers are represented above each curve. **D)** *E. coli* BACTH assay with T18-conjugated BacA constructs mutated in one (left) or both (right) polymerization interfaces, probed against the same constructs conjugated to T25, fusing each probe to either the N- or C-terminal of the BacA protein. Negative controls were performed against unfused T25 fragments. Positive controls (+) are shown on the bottom right of each plate.

In the N-to-N interface of the bactofilin domain, multiple residues have been shown to affect bactofilin polymerization in other species. Notably, *C. crescentus* V75 (*H. pylori* I55), which corresponds to V79 in *A. biprosthecum,* is important for *in vivo* and *in vitro* filament assembly (12,32). To test the importance of this residue in *A. biprosthecum,* we constructed a BacA_V79A_ mutant. We also mutated L46, predicted to lie at the N-to-N interface, whose substitution to a serine in *C. crescentus* showed fainter and larger _Cc_BacA_L46S_-mVenus foci and an increase in cell body fluorescence compared to WT (12). However, as serine is relatively small and only weakly hydrophilic, this substitution may not completely disrupt _Cc_BacA polymerization dynamics. Therefore, we constructed an L46R substitution. Additionally, we selected another conserved hydrophobic residue, I56, located at the N-to-N interface but on the opposite side of the L46 residue (Fig. 4B), creating an I56R substitution to test its role in polymerization.

In size-exclusion chromatography, the BacA_I56R_ mutant eluted mostly at around 9 mL, similar to purified WT BacA (Fig 4C [ii], dashed green line). On the other hand, BacA_L46R_ and BacA_V79A_ eluted mostly at 10-14 mL, which could correspond to smaller multimers of BacA, although BacA_L46R_ also showed a significant peak at around 9 mL, similar to WT BacA (Fig 4C [iii and iv], dashed orange and purple lines). Thus, these proteins carrying mutations in their N-to-N interfaces nonetheless appeared to retain some ability to polymerize. Consistently, these individual N-to-N interface mutants always retained some self-interaction in at least one configuration in a BACTH assay (Fig 4D), likely explained by their retention of the C-to-C polymerization interface.

To test mutations affecting both the N-to-N and C-to-C polymerization interfaces simultaneously, we created strains carrying the N-to-N interface substitutions in combination with the F134R substitution at the C-to-C interface. Mutating the N-to-N interface with either the I56R or L46R substitution in combination with the F134R substitution in the C-to-C interface resulted in a larger peak eluting at around 15 mL, corresponding to 22 kDa, the size of a BacA monomer (Fig. 4C [ii and iii], solid green and orange lines), consistent with a disruption of both polymerization interfaces. However, both BacA_I56R F134R_ and BacA_L46R F134R_ still showed a peak at around 9 mL, corresponding to the elution time of the WT BacA multimer. Nevertheless, in the BACTH assay, BacA_L46R F134R_ and BacA_I56R F134R_ did not display self-interaction (Fig 4D), suggesting that the multimers in their chromatography may be aggregates of misfolded proteins rather than true bactofilin polymers as with WT BacA, which strongly interacts with itself in the BACTH assay (Fig 1E and (29)). Similar aggregation was seen when the V79A substitution was made in combination with the F134R substitution; BacA_V79A F134R_ produced a multimeric peak at around 9 mL similar to BacA_L46R F134R_ and BacA_I56R F134R_ (Fig. 4C [iv], solid purple line).

Finally, we wanted to determine the effects of these polymerization mutations on BacA localization and stalk synthesis in *A. biprosthecum*. We examined this in *A. biprosthecum* using mVenus-tagged constructs of BacA carrying the different point mutations. Testing the expression of these mutant proteins in Western blots using anti-mVenus antibodies, we found that most of the mutations resulted in a reduced amount of BacA in the cell (Fig. S3). Fluorescence microscopy showed a diffuse cell body fluorescence for most of these mutants, as opposed to the discrete bilateral foci seen with WT BacA-mVenus (Fig. 5A). Nevertheless, BacA_L46R_-mVenus showed a few patchy foci near the stalk site, while BacA_I56R_-mVenus showed large, delocalized foci, suggesting that these mutations may impair BacA localization to a lesser extent than the other point mutations (Fig. 5A). In particular, BacA_I56R_-mVenus localization was reminiscent of BacA_ΔC_-mVenus localization, indicating that the I56R substitution may modify BacA filaments in a way that forms large bactofilin structures, which is in accordance with chromatography data from purified BacA_I56R_.

**Figure 5.**
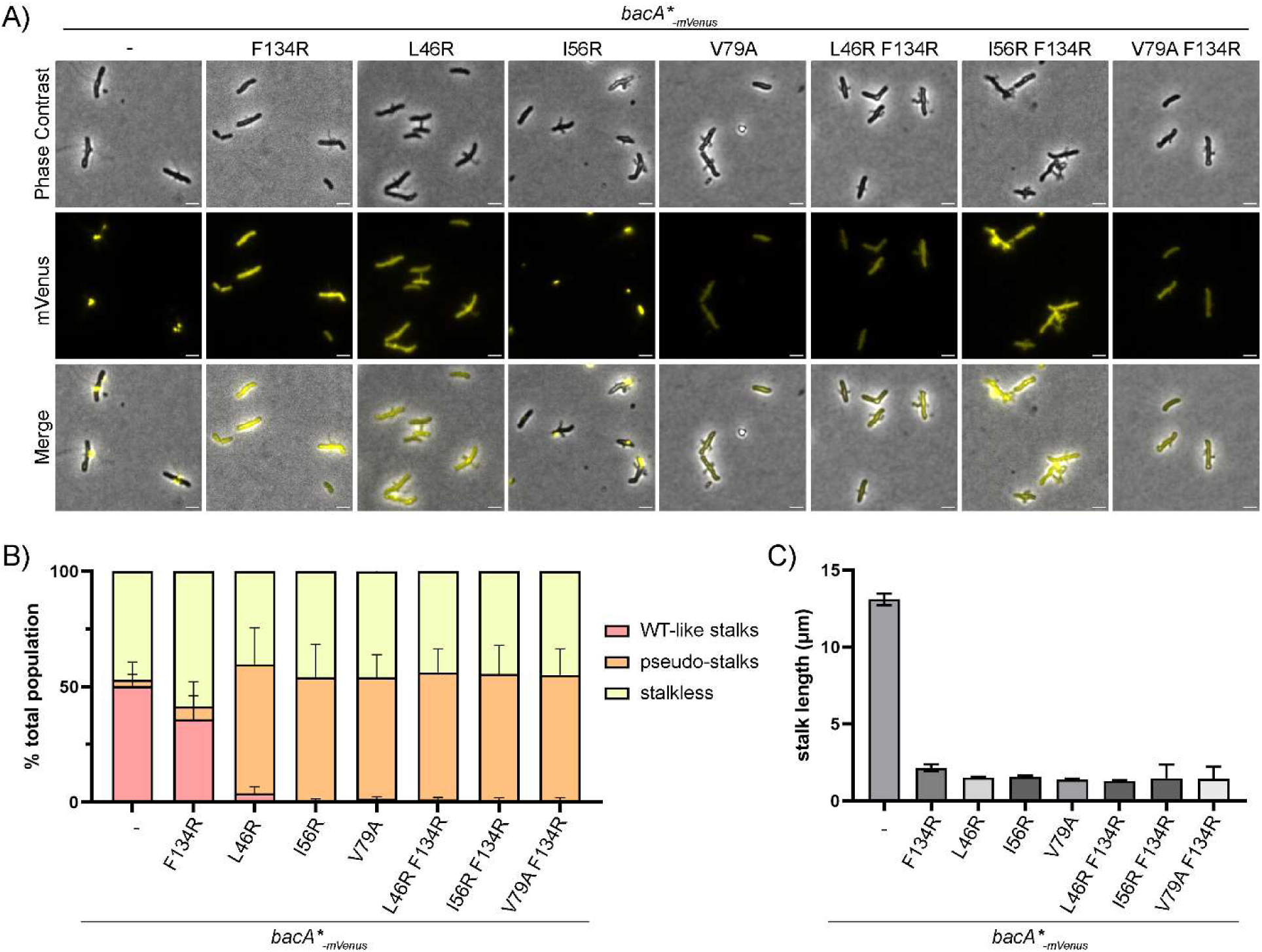
The BacA_F134R_ C-to-C interface polymerization mutant retains some functionality for stalk synthesis *in vivo* in *A. biprosthecum*. **A)** Phase contrast, fluorescence and merged microscopy images of BacA-mVenus strains carrying various polymerization mutations, grown in phosphate-limited HIGG medium. For better visualization, LUTs were adjusted to reduce the brightness of the BacA-mVenus and BacA I56R-mVenus fluorescence signals, as their punctate localization showed much higher intensity compared to the diffuse signals in other strains. See Figure S3B for fluorescent images of these two mutants with identical LUTs.Scale bars = 2 μm. **B)** Percentage of cells with different types of stalks for each of the BacA-mVenus polymerization mutant strains. Based on phase contrast images from three independent biological replicates, stalks were assessed and categorized either as WT-like stalks or pseudostalks. Data are presented as the mean percentage of cells with WT-like stalks or pseudostalks. Error bars represent SD. Total numbers of cells for each strain were as follows: WT n=1940; F134R n=1312; L46R n=1817, I56R n=1645; V79A n=1483; L46R F134R n=1308; I56R F134R n=1907; V79A F134R n=1746. **C)** Distribution of stalk/pseudostalk lengths in BacA-mVenus polymerization mutant strains. Error bars represent SD. Total numbers of cells for each strain were as follows: WT n=189; F134R n=63; L46R n=192, I56R n=113; V79A n=167; L46R F134R n=155; I56R F134R n=38; V79A F134R n=70)

We also used phase contrast microscopy of the different polymerization mutants to assess stalk formation and stalk length. Given that the C-to-C polymerization interface mutant BacA_F134R_ lacks the ability to polymerize, we expected it to lead to impaired stalk formation. But surprisingly, we observed stalks in 41% of BacA_F134R_ cells (compared to 50% in WT) (Fig 5B). Most of the stalks were short and thin (Fig. 5A), meaning that impairment of C-to-C polymerization does not completely inhibit stalk formation, but restricts them in terms of length (Fig 5C). In contrast, all N-to-N interface mutants had pseudostalks, both by themselves and in combination with the F134R mutation. In these strains, pseudostalks were observed in a similar proportion of cells as stalked cells in the WT background (Fig 5A-B). Their pseudostalks were highly reminiscent of the *ΔbacA* phenotype, similar in proportions and stalk lengths, and resulting in the same sorts of cell abnormalities as previously described for *ΔbacA* (29).

## Discussion

Over the past few decades, it has become evident that filament-forming cytoskeletal proteins are important for bacterial cellular organization (1,2). Among these, bactofilins, a relatively recently discovered cytoskeletal filament, are highly conserved, widely distributed, and implicated in a variety of cellular processes including morphology (7,9,14,15,22,31). The bactofilin proteins studied thus far share the common characteristics of 1) a conserved β-helical core (the bactofilin domain) flanked by variable N- and C-terminal regions and 2) the ability to self-polymerize without requiring cofactors. Using *A. biprosthecum* as a model system, we were able to show that these disordered N- and C-terminal regions are important for BacA’s role in stalk synthesis. Moreover, we showed that BacA polymerization is also important for its role in stalk synthesis, relying on critical residues at both the N-to-N and C-to-C polymerization interfaces.

In many organisms, bactofilins act as a morphogenesis protein through interactions with peptidoglycan-associated proteins. Often, they are found in the same operon as a peptidoglycan hydrolase LmdC, which has been found to be important for morphogenesis in *H. neptunium* and *R. rubrum* (23). In *H.* pylori, the bactofilin CcmA interacts with various Csd (cell shape determining) proteins. In *C. crescentus* the bactofilins BacA and BacB interact with the PG-synthetic protein PbpC (9,15). The presence of proline-rich domains in both the N- and C-terminal domains of bactofilins may facilitate their versatile interactions with other morphogenic proteins, potentially allowing them to regulate peptidoglycan synthesis akin to what has been observed with other peptidoglycan-related scaffolding proteins.

In recent years, the N-terminal domains of bactofilins have been implicated in membrane binding. In a limited subset of Gammaproteobacteria, bactofilins even have transmembrane domains within their N-terminal regions, although most bacteria instead rely on a short amphipathic peptide sequence for membrane associations (8,34,35). In our study, we observed that deleting the N-terminal region of BacA led to a pseudostalk phenotype as severe as observed with deletion of the entire *bacA* gene. Using an mVenus-tagged fusion, we found that BacA_ΔN_ was still able to form foci *in vivo,* but we found that these foci were mislocalized throughout the cell. This is in contrast to what has been found in *C. crescentus*, where mutations in the N-terminal membrane-binding region, result in a diffuse localization of _Cc_BacA underscoring a potential interdependence between polymerization and membrane binding (35). It is possible that this difference is due to the different recruitment role played by SpmX in the *Asticcacaulis* genus (25).

In *A. biprosthecum*, SpmX is responsible for the recruitment of the stalk synthesis machinery, including BacA, to the base of the stalk. While interactions between SpmX and BacA_ΔN_ were not lost in a BACTH assay, SpmX nonetheless does not seem to be able to recruit BacA_ΔN_ to the base of the stalk, as seen by the general absence of BacA_ΔN_-mVenus foci at the stalk base, and its pseudostalk phenotype reminiscent of a *ΔbacA* mutant (29). Meanwhile, deletion of the N-terminal domain of BacA had no detectable effect on SpmX localization, even though BacA_ΔN_ is not localized at the stalk base. Based on these results, we hypothesize that the N-terminal domain of BacA, perhaps through its interactions with the membrane, plays a role in its recruitment to the base of the stalk. Loss of the N terminal domain of BacA thereby prevents its localization at the stalk base, leading to a pseudostalk phenotype.

Shifting focus to the C-terminal domain of bactofilins, previous studies suggest that it plays an important role in protein-protein interactions involving the bactofilins. In *M. xanthus*, the C-terminal domains of its bactofilins interact with the PadC-ParA chromosome segregation complex (19). In this species, fluorescent tagging on the C-terminus of the bactofilins failed to give proper localization or complement gene deletion, likely due to molecular crowding from the fluorescent proteins preventing its protein-protein interactions (18,19). Similarly, in *P. mirabilis*, a transposon insertion at the C-terminus of the bactofilin CcmA caused a more severe morphological phenotype than gene deletion, indicating that such an insertion disrupts other functions in the cell (16). These observations underscore the crucial role of the C-terminal domain in bactofilin activity, particularly for protein-protein interactions. In line with these findings, the C-terminal domains of the bactofilins BacA in *C. crescentus* interact with PG synthetic proteins such as PbpC in addition to the core domain (35). PbpC lacks homologs in the *Asticcacaulis* genus, and the specific PG synthetic enzymes required for stalk synthesis have not yet been identified in *A. biprosthecum*. Here, we determined that the BacA C-terminal domain is required for interactions with SpmX, which in turn is important for BacA localization to the stalk synthesis site. In the absence of the C-terminal domain, BacA_ΔC_-mVenus mislocalizes, preventing its recruitment to the base of the stalks as discrete foci, in contrast WT BacA-mVenus. Interestingly, BacA_ΔC_ cells produce multiple phenotypes ranging from WT-like stalks to Δ*bacA*-like pseudostalks. Specifically, many cells show a widened stalk base extruding thin stalks at their tips. This phenotype is associated with elongated BacA_ΔC_-mVenus foci at the base of these thin stalks (either at the regular stalk base or within the larger pseudostalk-like base of the extended thin stalk). This suggests that despite the absence of the C terminal domain and its interactions with SpmX, BacA_ΔC_ can nonetheless interact with other stalk synthesis proteins recruited by SpmX at the stalk base, allowing rescue of WT-like stalk synthesis after an initial round of peptidoglycan insertion in a pseudostalk-like manner. This suggests that (1) the SpmX-BacA interaction is not required throughout the entire stalk synthesis process and (2) the C-terminal domain is not the only domain mediating BacA’s protein-protein interactions for stalk synthesis, which may also be mediated through the core bactofilin domain or the N-terminal domain, as observed in *H. pylori* CcmA-Csd protein interactions (15,20,32).

Bactofilins have been characterized by the presence of the DUF583 domain, i.e. the bactofilin domain, comprising a β-helical structure with a hydrophobic core (8,12). Bactofilins polymerize via their hydrophobic domain interactions to form filaments and *in vivo* structures. The critical role of bactofilin polymerization is supported by findings in other species, where polymerization disruptions lead to significant functional impairments. Here, we mutated specific hydrophobic residues predicted to mediate interactions at the N-to-N and C-to-C interfaces to test the importance of BacA polymerization for its function in *A. biprosthecum*.

As reported previously, we confirmed that BacA forms non-polar filaments based on BACTH and size-exclusion data. We were also able to construct a fully monomeric bactofilin by mutating both the N-to-N and C-to-C polymerization interfaces with the L46R and F134R mutations. When we examined the effect on stalk synthesis of amino acid substitutions that disrupt polymerization, we expected non-polymerizing mutants to completely phenocopy the *ΔbacA* mutant, i.e. that they would produce pseudostalks irrespective of the mutated interface, since they should not be able to form higher-order structures that allow bactofilin function as a stalk-specific scaffolding protein. Interestingly, while the F134R mutation on the C-to-C interface leads to dimer formation *in vitro* and diffuse localization *in vivo*, stalk synthesis was not completely abolished in this strain. Instead, BacA_F134R_ cells appear to produce mostly WT-like stalks, albeit shorter in overall length. Out of all the polymerization mutants we tested, the F134R mutation on the C-to-C interface was the only one showing this phenotype *in vivo,* whereas all other polymerization mutants, which targeted the N-to-N interface, demonstrated pseudostalk phenotypes similar to *ΔbacA*. Our hypothesis is that BacA_F134R_ dimers may be sufficient for initiation of stalk synthesis, allowing the formation of short WT-like stalks. However, BacA polymerization is needed to stabilize or coordinate extended PG synthesis at the base of the stalk for efficient stalk elongation.

This hypothesis is in accordance with the observation that _Cc_BacA-PbpC interaction is dependent on _Cc_BacA polymerization (35) and the fact that BacA/B deletions produce shorter stalks in *C. crescentus* but do not abolish stalk synthesis (9). Similarly, in the stalked budding bacterium *H. neptunium*, the F130R mutant forms partial stalks but fails to generate budding daughter cells (23).

Our study underscores the importance of both polymerization and the N- and C-terminal domains of BacA in *A. biprosthecum* stalk synthesis. Our results suggest that while the conserved β-helical core domain is vital for polymerization and functionality, the flanking domains may contribute to species-specific functions, reflecting modular evolution. The terminal domains of the bactofilins vary greatly in sequence and length among species. Considering the diverse mechanisms through which bactofilins appear to be acting and interacting, it is conceivable that they serve as a versatile platform for the generation of novel functions through terminal domain modifications to achieve different scaffolding properties and regulatory interactions.

## Methods

### Bacterial strains and growth conditions

All freezer stocks were maintained in 10% DMSO at -70°C*. A. biprosthecum* strains used in this study were grown at 26°C in Peptone Yeast Extract (PYE) medium on plates or in liquid culture, supplemented with 5 μg ml^-1^ kanamycin where appropriate. For microscopy of cells grown in low-phosphate <u>H</u>utner base-imidazole-buffered-<u>g</u>lucose-<u>g</u>lutamate (HIGG) medium (36), strains from PYE liquid cultures were washed 2X with double distilled H_2_O before being added in a 1:20 ratio in low-phosphate HIGG, and grown at 26°C with shaking for 48h before imaging. Cells for imaging SpmX localization were grown in HIGG for 24 hours prior to microscopy.

A) *E. coli* strains used in this study were grown in liquid lysogeny <u>b</u>roth (LB) medium at 37°C (or 30°C for BACTH assays) supplemented with antibiotics or supplements as necessary (carbenicillin 100 µg ml^-1^, kanamycin 50 µg ml^-1^, X-gal (5-Bromo-4-chloro-3-indolyl β-D5-Bromo-4-chloro-3-indolyl β-D-galactopyranoside-galactopyranoside) 40 µg ml^-1^, 0.3 mM diaminopimelic acid and 0.5 mM IPTG).

Mating and electroporation of *A. biprosthecum* were performed as previously described (29). Outgrowth was performed for 8-24h at 26°C before plating. Mating of plasmids into *A. biprosthecum* was performed using the *dap*^-^ *E. coli* strain WM3064 (YB7351) (37). Allelic exchange and deletions were achieved using a two-step sucrose (3% final) counter selection procedure with pNPTS138. In-house stocks of chemically competent DAP auxotroph (YB7351) and BTH101 (YB9171) *E. coli* cells were prepared as previously described (29). Plasmids were introduced into *E. coli* strains using chemical transformation according to the manufacturer’s protocols. All strains used in this study can be found in Table S1 and S2.

### Recombinant DNA methods

DNA amplification, Gibson cloning, and restriction digests were performed according to the manufacturers’ instructions, using iProof High-Fidelity DNA Polymerase (Bio-Rad Laboratories, Cat. No. 1725302) and NEBuilder HiFi DNA Assembly Master Mix (New England Biolabs, Cat. No. E2621X). All primers for Gibson assembly were designed using NEBuilder Assembly Tool (New England Biolabs) and are available upon request. A detailed list of plasmids can be found in Table S2. Restriction enzymes and the Gibson cloning mix were purchased from New England Biolabs. Cloning steps were carried out in *E. coli* (alpha-select competent cells, Bioline) and plasmids were purified using GeneJET Plasmid Miniprep Kit (ThermoFisher). Sanger sequencing was performed by the Indiana Molecular Biology Institute (Indiana University) or the IRIC sequencing platform (Université de Montreal) with double stranded plasmid or PCR templates, which were purified with a GeneJet PCR purification kit (ThermoFisher). Whole plasmid sequencing was performed by Plasmidsaurus using Oxford Nanopore Technology with custom analysis and annotation.

### BACTH interaction assays

pUT/pUTc18 fusion and pKT/pKNT25 fusion plasmids containing *bacA,* its mutant constructs, or *spmX* were transformed into BTH101 *E. coli* cells and selected on LB plates with carbenicillin and kanamycin. Colonies were grown overnight in liquid LB with antibiotics and spotted on plates containing X-Gal and IPTG. After overnight incubation at 30°C, plates were imaged using a smartphone camera.

### Light microscopy and fluorescence imaging

For light microscopy analysis, cells were spotted onto pads made of 1% SeaKem LE Agarose (Lonza, Cat. No. 50000) in HIGG and topped with a glass coverslip. Images were recorded with inverted Nikon Ti-E or Ti-2 microscopes with CFP/YFP/mCherry filter cubes using either 1) a Plan Apo 60X 1.40 NA oil Ph3 DM objective and an iXon X3 DU885 EMCCD camera or 2) a Plan Apo λ 100X 1.45 NA oil Ph3 DM objective and either a Photometrics Prime 95B sCMOS camera or an Andor iXon3 DU885 EM CCD camera.

### Image analysis

Stalk length and stalk percentage data were obtained using FIJI (<u>F</u>iji Is <u>J</u>ust ImageJ). Briefly, phase micrographs were imported into FIJI and stalks were manually traced using the Freehand Line tool and stalk lengths were determined from the manual trace using the Measure function. Data were then plotted on a log10 scale on the y-axis and represented as box and whisker plots where the middle line represents the median, the lower and upper hinges correspond to the first and third quartiles (the 25th and 75th percentiles) and outliers outside 1-99% are plotted individually. A mirrored-density violin plot is underlaid to show the continuous distribution of the data. Percentages of stalked cells per image were calculated by manually counting the number of stalked and non-stalked cells per image. Stalk width measurements were made on AFM height images using Nanoscope software (Bruker, Santa Barbara, CA).

Subcellular localization heatmaps were generated using the ImageJ plugin MicrobeJ (version 5.13o/p) (38). Cell silhouettes and cell body outlines were produced by importing phase micrographs into Illustrator (Adobe Inc.) and manually tracing the silhouette or outline.

### Bioinformatic analyses and AlphaFold predictions

Genomic and protein sequence data were obtained from the Integrated Microbial Genomes (IMG) database (39). Multiple sequence alignment of bactofilin sequences was performed using Jalview (40) . Disorder plots were generated using SPOT-Disorder2 (41), IUPred 3 (42), Md2 (43), and Espritz (44). All were normalized and combined to assemble a single graph to predict BacA’s disordered domains. Structures of the BacA monomer and tetramer were predicted using AlphaFold-multimer as implemented through ColabFold (45). All predicted structures for the BacA monomer or tetramer can be found in Figure S4, along with their predicted aligned error scores. Rank 1 predictions were used. For the BacA tetramer, only the core bactofilin domain was used for prediction. Protein structures were visualized and superimposed using ChimeraX (46).

### Protein purification

Recombinant plasmids overproducing His6-BacA with or without mutations were transformed into the BL21 *E. coli* strain. Transformants were grown at 37°C until OD_600_ = 0.6 in LB medium with kanamycin (50 μg ml^−1^). His6-BacA protein expression was induced for 3h at 30°C by adding 0.5 mM IPTG (isopropyl β-D-thiogalactopyranoside). Cells were centrifuged, and pellets were resuspended in 1/25 volume of Buffer A (50 mM HEPES at pH 8, 300 mM NaCl, 2 mM β-mercaptoethanol) and lysed by sonication (20 sec ON / 40 sec Off, 5 min, Misonix S4000). Cell debris was pelleted by centrifugation at 36,000 g for 30 min at 4°C. The supernatant was loaded onto a Ni-NTA resin (Qiagen) column in an AKTA FPLC pure system. After washing with Buffer A, His6-BacA protein was eluted with a linear gradient of Buffer B (50 mM HEPES at pH 8, 300 mM NaCl, 2 mM β-mercaptoethanol, 500 mM imidazole). Elution fractions were loaded onto SDS page gels and the peak fractions containing the His6-BacA protein were pooled.

Upon purification, His6-tagged WT and mutant BacA proteins were used for a polymerization assay performed by dialyzing the purified protein in Buffer C (25 mM HEPES at pH 8, 250 mM NaCl, 2 mM β-mercaptoethanol). Purified proteins were used for AFM, electron microscopy, or phase contrast microscopy and size-exclusion chromatography.

### Size-exclusion chromatography

500 µl of purified proteins from the polymerization assay were loaded onto a Superdex 200 Increase 10/300 GL column (GE Healthcare) using an AKTA FPLC pure system. Protein elution was monitored using UV and normalized to 100% of the highest observed elution. The column was calibrated with standard protein controls to correlate elution time/volume to protein molecular weight.

### AFM microscopy

10 µl of cell suspensions (for whole cell imaging) or diluted proteins (for filament formation) were applied to glass coverslips and left to dry for 2 hours. The dried cells or proteins were imaged using a Bioscope Resolve AFM (Bruker, Santa Barbara, CA) atomic force microscope using contact mode in air.

### Electron microscopy

10 µl of cell suspension was applied to an electron microscopy grid (Formvar/Carbon on 300 mesh; Ted Pella Inc., Cat. No. 01753-F) for 5 min at room temperature. Excess liquid was removed with Whatman filter paper. Cells were then negatively stained with 10 µl 1% uranyl acetate (UA) and excess UA liquid was immediately removed with Whatman filter paper. Grids were allowed to dry, stored in a grid holder in a desiccation chamber, and imaged with a kV JEOL JEM 1010 transmission electron microscope (JEOL USA Inc.).

### Western blot analysis

All strains were grown until OD = 1.0 before being centrifuged and resuspended in 50 μl of 50mM Tris, 200 mM NaCl supplemented with 0.1 μl of Universal Nuclease (Pierce #88700) added with Laemmli buffer. 15 μl of each sample was loaded onto 4%–20% precast polyacrylamide gels (BioRad) before being transferred to a nitrocellulose membrane according to the manufacturer’s instructions. Loading was controlled by Ponceau’s staining before immunoblotting was performed by addition of the appropriate primary antibody, either an anti-GFP (anti-mVenus) polyclonal antibody (MBL #598), a custom made anti-BacA polyclonal antibody, or an anti-GAPDH polyclonal antibody (Proteogenix, France). Goat anti-rabbit HRP (Pierce) was used as the secondary antibody. Transferred blots were visualized with SuperSignal West Pico substrate (ThermoFisher Scientific) using a Bio-Rad Chemidoc.

## Acknowledgments

Early stages of this work were supported by NIH grant R35GM122556 to Y.V.B. Current work in the Y.V.B. lab is supported by a Canada 150 Research Chair in Bacterial Cell Biology. We thank Vaidehi Patel for the SpmX-mCherry plasmid, previous and current Brun Lab members for advice and scientific discussions, and Velocity Hughes (Synthesis by Velocity, Malmö, Sweden) for editorial help with the manuscript.

## Author Contributions

M.J., P.D.C., and Y.V.B. designed the research and wrote the manuscript. M.J. and P.D.C. performed experiments and analyzed data. Y.V.B. acquired funding.

## Competing interests

The authors declare no competing interests.

---

**Figure S1.**
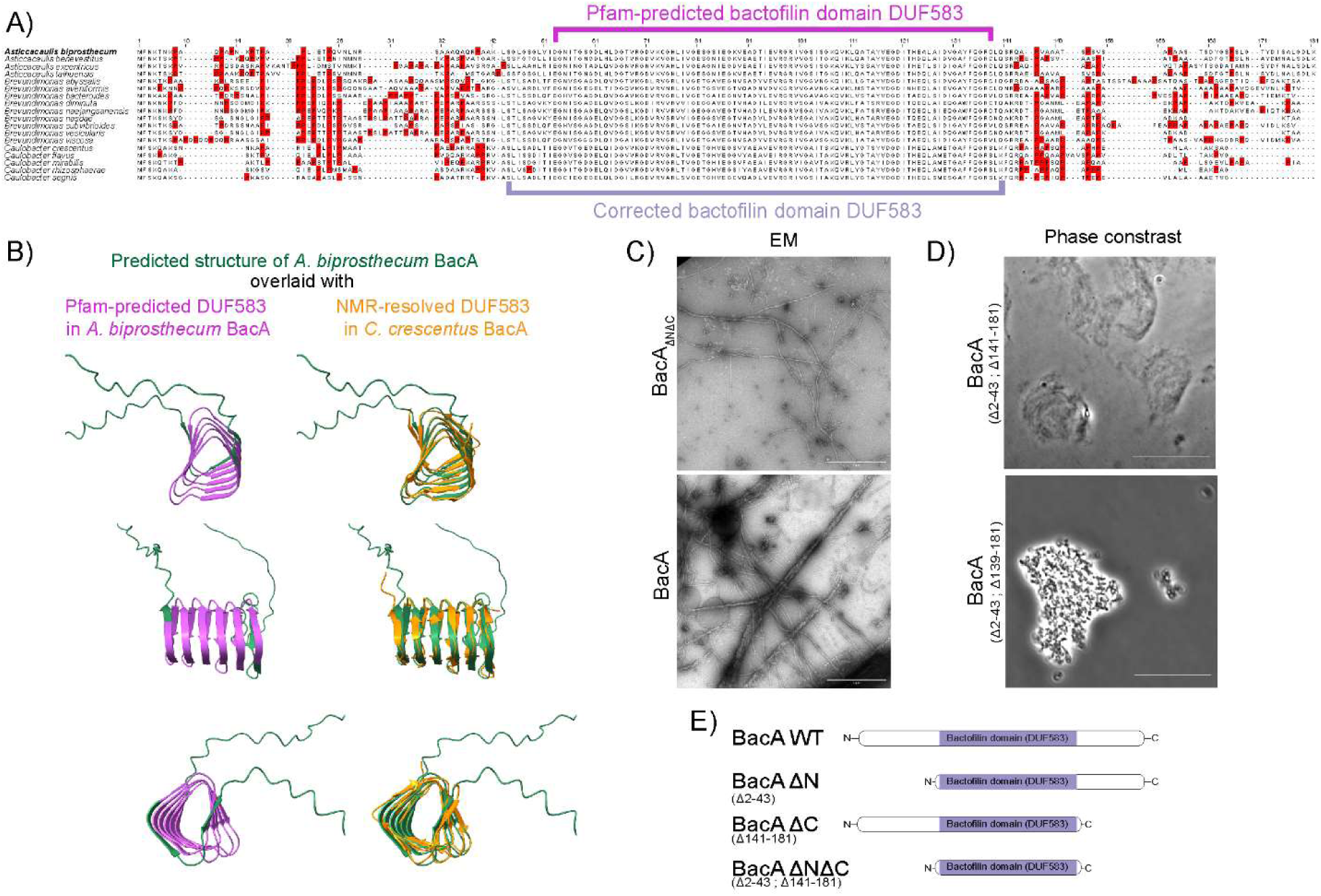
Analysis of the domain structure of bactofilins in the *Caulobacteraceae* family, and proposed *A. biprosthecum* BacA mutations. **A**) Multiple sequence alignment from Figure 1A highlighting proline residues (highlighted in red) in the proline-rich N- and C-terminal domains flanking the central bactofilin domain (DUF583). The Pfam-predicted bactofilin domain is delineated in magenta and the corrected bactofilin domain in mauve. **B**) Left: The predicted structure of *A. biprosthecum* BacA superimposed with the NMR-resolved structure of *C. crescentus* BacA (orange) (PDB-ID : 2N3D). Right: Structural prediction of *A. biprosthecum* BacA (green) using AlphaFold, with the Pfam-delimited bactofilin domain (DUF583) in magenta. Side, top, and bottom views of the structures are presented. **C**) High-resolution EM images of purified BacA and BacA_ΔNΔC_ filaments (scale bar = 1 µm). **D**) Phase contrast images of purified BacA_Δ2-43;Δ141-181_ (BacA_ΔNΔC_) filaments and BacA_Δ2-43;Δ139-181_ aggregates (scale bar = 20 µm). **E**) Schematic of the full-length BacA protein from *A. biprosthecum* and the proposed mutants for this study, based on the corrected bactofilin domain (residues 44-140).

**Figure S2.**
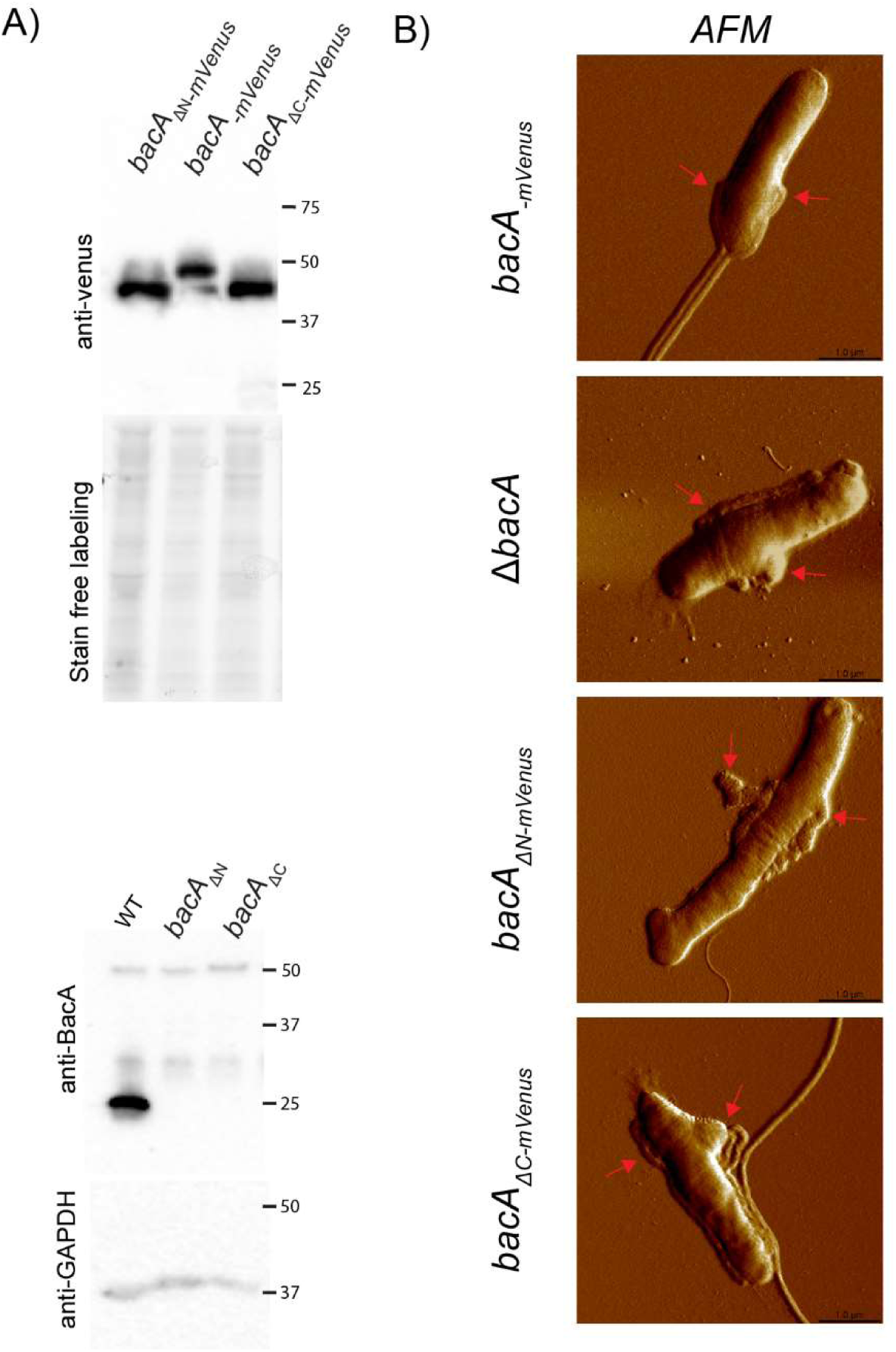
Protein expression and AFM microscopy of BacA mutants strains. **A**) Top: Western blots using anti-GFP antibodies in mVenus-tagged BacA terminal domain deletion strains in *A. biprosthecum*. Strain free labeling was used as a loading control. Bottom: Western blots using anti-BacA antibodies in untagged terminal domain deletion strains in *A. biprosthecum* showed that antibodies raised against full length BacA protein were unable to detect truncated BacA mutants. Anti-GAPDH antibodies were used as a loading control. **B**) Atomic force microscopy of *A. biprosthecum bacA-mVenus*, Δ*bacA*, *bacA*_Δ_*N-mVenus* and *bacA*_Δ_*C-mVenus* strains, used to analyze the width of the stalk base. Stalks/pseudostalks are indicated with red arrows. Cells were grown in phosphate-limited (HIGG) medium (see Methods). Scale bars = 1 μm.

**Figure S3.**
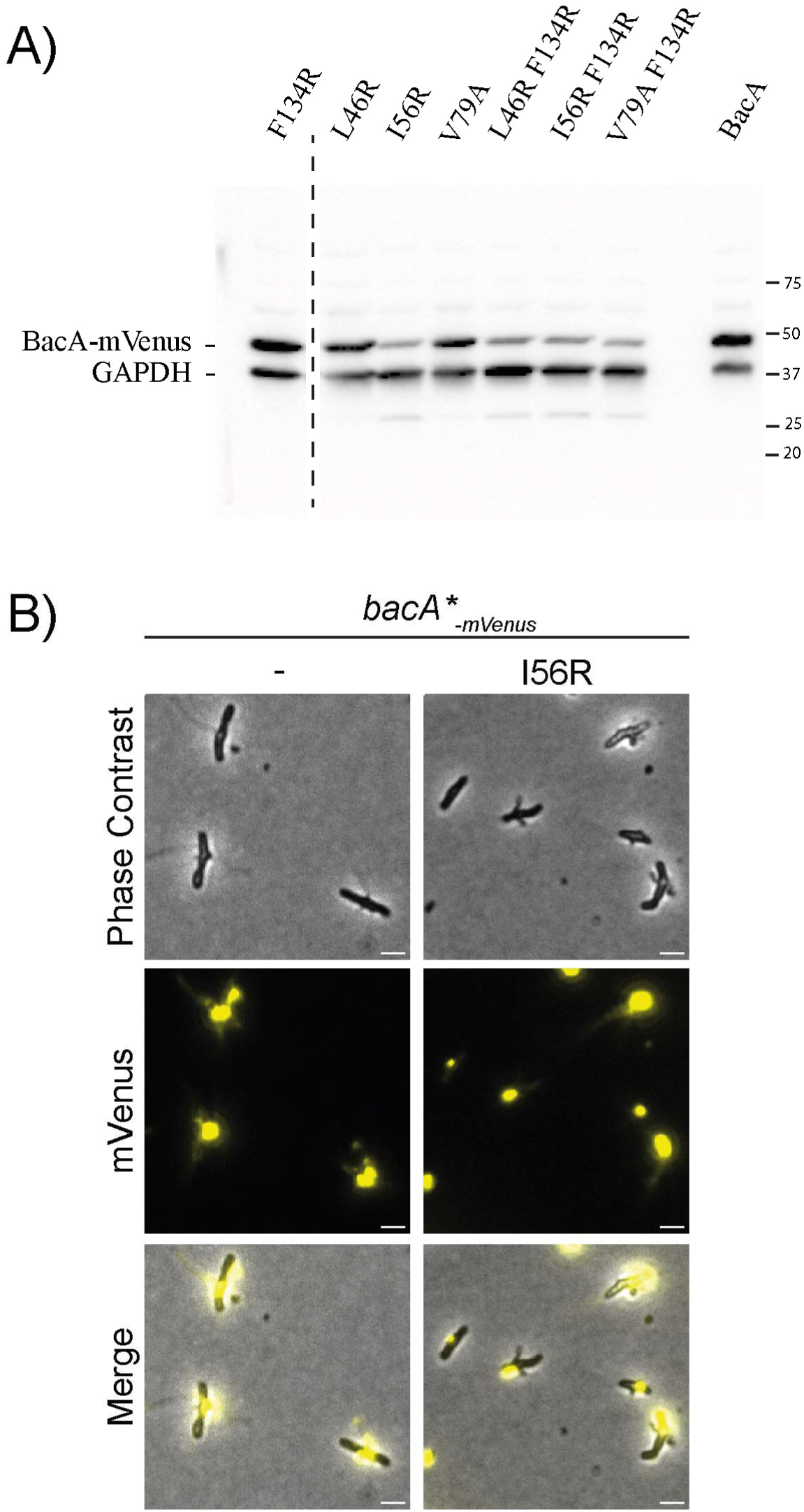
Western blot to verify the expression of the BacA polymerization point mutants. **A)** Western blot using anti-GFP antibodies in the BacA polymerization point mutants in *A. biprosthecum*. Cells lysates were loaded at the same level of protein. Anti-GAPDH is presented as a loading control. **B)** Phase-contrast, fluorescence, and merged microscopy images of BacA-mVenus and BacA I56R-mVenus, with LUTs matching those used for mutants displaying diffuse fluorescence presented in Figure 5A.

**Figure S4:**
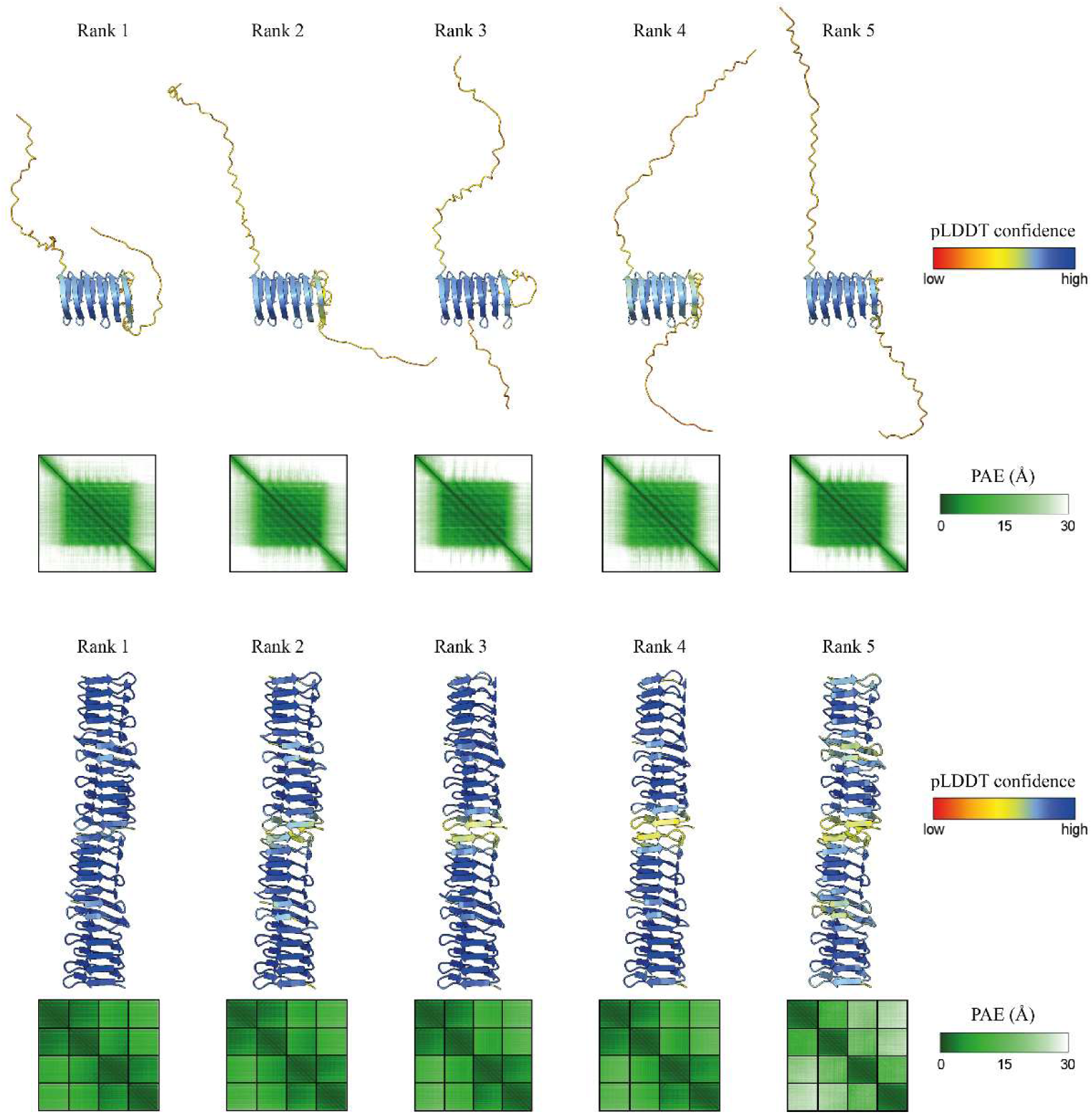
BacA monomer and tetramer structural predictions. BacA monomer (top) and tetramer (bottom) predicted by AlphaFold, used in Figure 1 and 4 respectively. Structures are aligned and shown from the same orientation. Structures are ranked according to the predicted template modeling (pTM) score and are colored according to the predicted local distance difference test (pLDDT) score. Confidence in the prediction of each complex is indicated by the predicted aligned error (PAE) scores, which indicate positional error in angstroms for a given pair of residues across both protein chains.

**Table S1.** Strains used in this study.

| Strain (YB#) | Genotype/description | Construction | Reference/source |
| --- | --- | --- | --- |
| *E. coli* |  |  |  |
| 105 | Overexpression strain; BL21λDE3 (F- ompT hsdSB (rB-mB-) gal dcm (IDE3) |  | - |
| 7351 | DAP auxotroph; WM3064 (thrB1004 pro thi rpsL hsdS lacZΔM15 RP4-1360 Δ(araBAD)567 ΔdapA1341::[erm pir]) |  | (37) |
| 9171 | BACTH strain; BTH101 (F-, cya-99, araD139, galE15, galK16, rpsL1 (Str r), hsdR2, mcrA1, mcrB1.) |  | Euromedex EUB001 |
| *A. biprosthhecum* |  |  |  |
| 642 | WT C19 |  | (26) |
| 8597 | ΔbacA | - | (29) |
| 9139 | *bacA* (aa 45-181)-*mVenus* (ΔN) | Created by electroporating and double recombination using pMJ77 into *A. biprosthhecum* C19. | This study |
| 9141 | *bacA-mVenus* | Created by electroporating and double recombination using pMJ79 into *A. biprosthhecum* C19. | This study |
| 9189 | *bacA*(aa 1-140)-*mVenus* (ΔC) | Created by electroporating and double recombination using pMJ76 into *A. biprosthhecum* C19. | This study |
| 9506 | *bacA F134R-mVenus* | Created by electroporating and double recombination using pMJ125 into *A. biprosthhecum* C19. | This study |
| 9507 | *bacA L46R-mVenus* | Created by electroporating and double recombination using pMJ124 into *A. biprosthhecum* C19. | This study |
| 9510 | *bacA I56R-mVenus* | Created by electroporating and double recombination using pMJ123 into *A. biprosthhecum* C19. | This study |
| 9512 | *bacA V79A mVenus* | Created by electroporating and double recombination using pMJ159 into *A. biprosthhecum* C19. | This study |
| 9513 | *bacA V79A F134R-mVenus* | Created by electroporating and double recombination using pMJ156 into *A. biprosthhecum* C19. | This study |
| 9594 | *bacA L46A F134R-mVenus* | Created by electroporating and double recombination using pMJ127 into *A. biprosthhecum* C19. | This study |
| 9595 | *bacA I56A F134R-mVenus* | Created by electroporating and double recombination using pMJ128 into *A. biprosthhecum* C19. | This study |
| **9895** | bacA (aa 1-140) (ΔC) | Created by electroporating and double recombination using pMJ169 into *A. biprosthhecum* C19. | This study |
| **9896** | bacA (aa 45-181) (ΔN) | Created by electroporating and double recombination using pMJ170 into *A. biprosthhecum* C19. | This study |
| **10188** | bacA (aa 1-140) (ΔC); spmX-mCherry | Created by electroporating and double recombination using pNPTS138-SpmX-mCherry into *A. biprosthhecum* YB9895. | This study |
| **10189** | bacA (aa 45-181) (ΔN); spmX-mCherry | Created by electroporating and double recombination using pNPTS138-SpmX-mCherry into *A. biprosthhecum* YB9896. | This study |
| **10190** | spmX-mCherry | Created by electroporating and double recombination using pNPTS138-SpmX-mCherry into *A. biprosthhecum* C19. | This study |

**Table S2.**
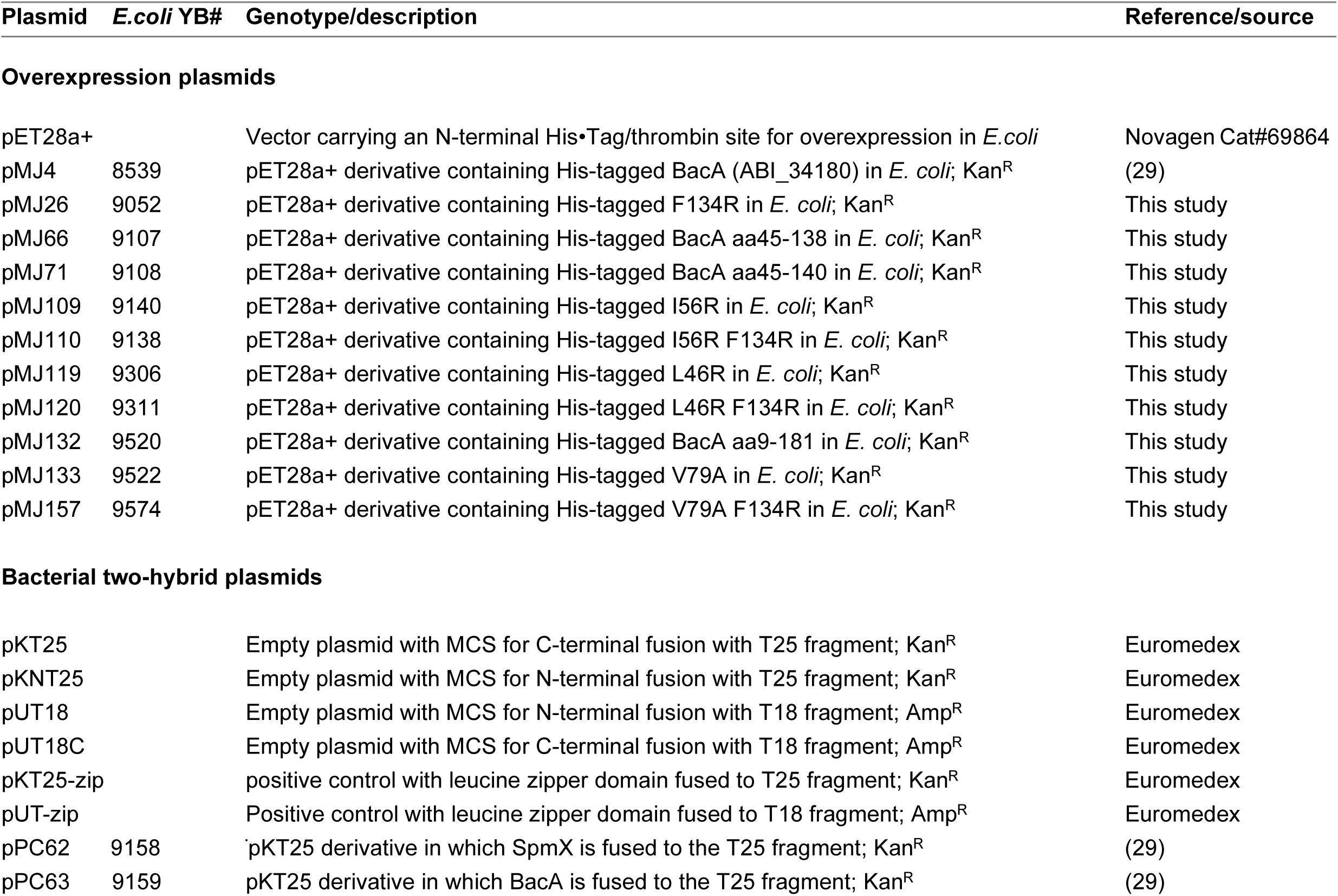
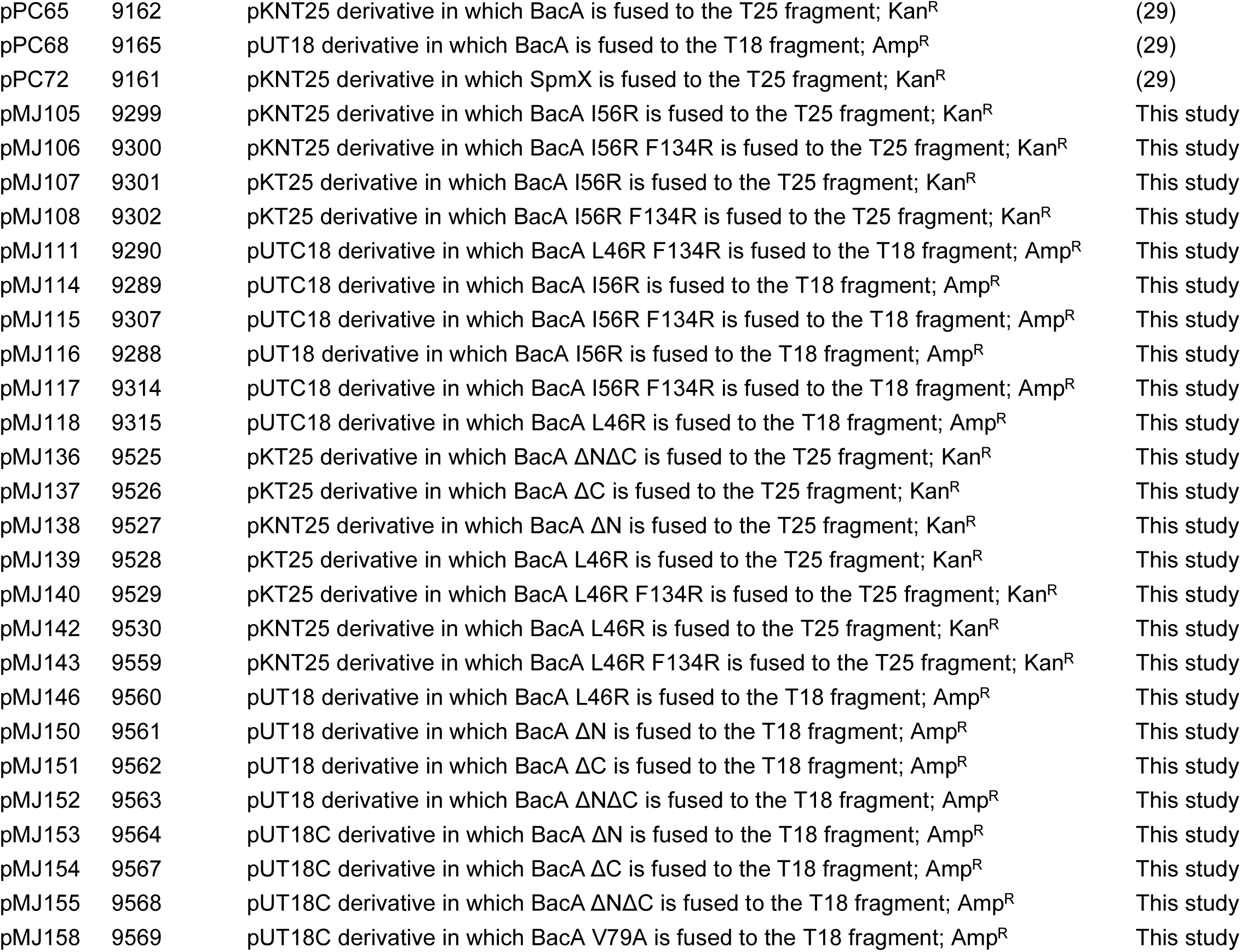
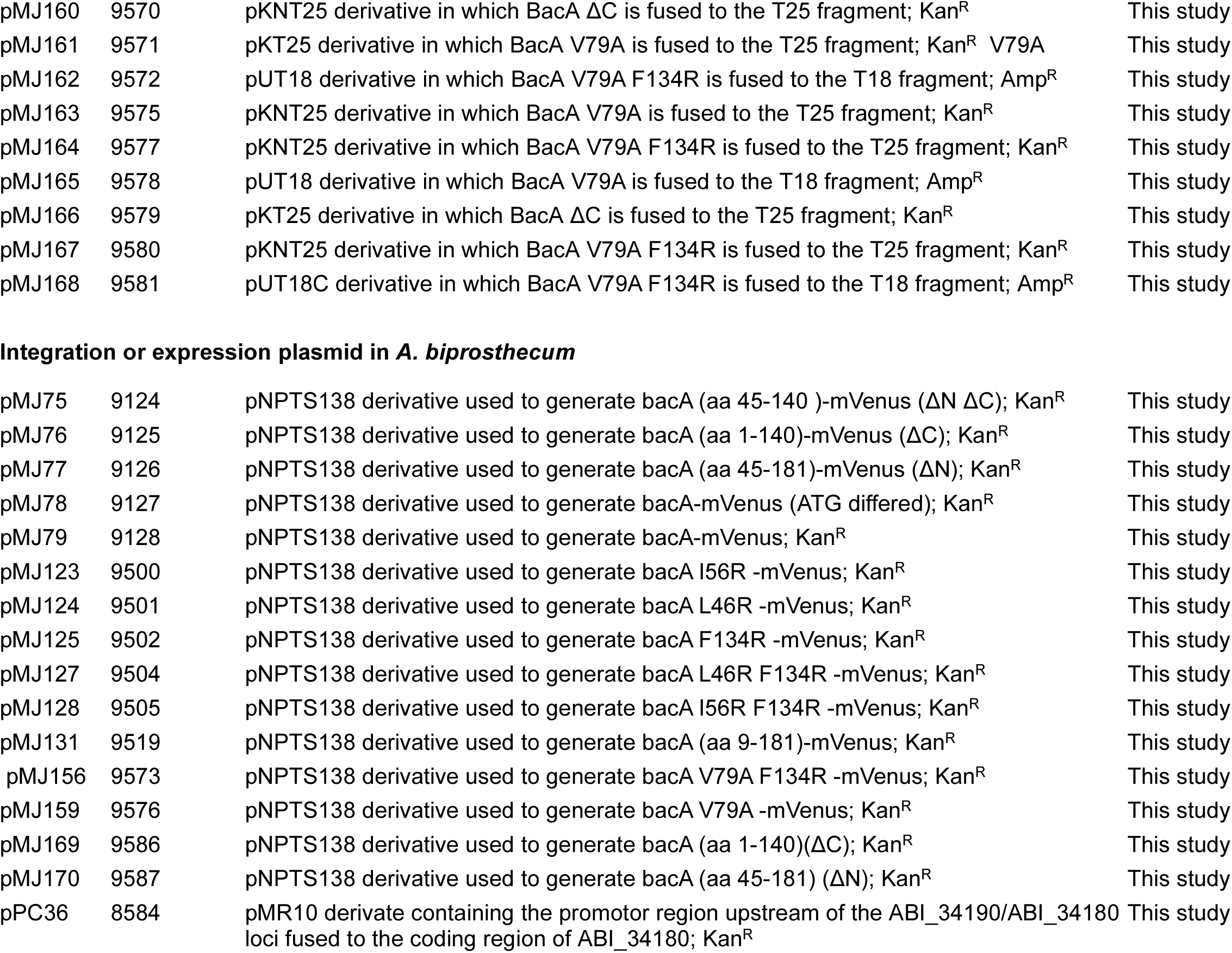
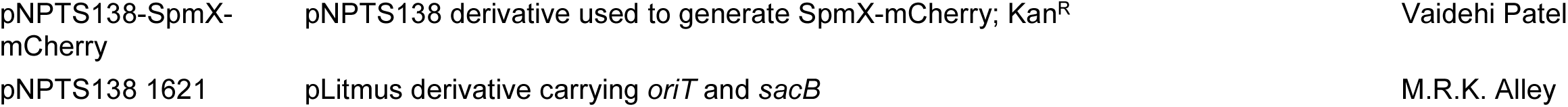
Plasmids used in this study.

## Notes

### Competing Interest Statement

The authors have declared no competing interest.

